# Nearshore corals on the Mesoamerican Barrier Reef System on pace to cease growing as soon as year 2110

**DOI:** 10.1101/298158

**Authors:** Justin H. Baumann, Justin B. Ries, John P. Rippe, Travis A. Courtney, Hannah E. Aichelman, Isaac Westfield, Karl D. Castillo

## Abstract

Anthropogenic global change and local anthropogenic stressors are decreasing coral growth and survival globally, thus altering the structure and function of coral reef ecosystems. We show that skeletal extension rates of nearshore colonies of *Siderastrea siderea* and *Pseudodiploria strigosa* across the Belize Mesoamerican Barrier Reef System (MBRS) have declined at average rates of 0.01 and 0.08 mm/yr, respectively, over approximately the past century, while offshore conspecifics exhibited no significant trend in extension with time. This caused extension rates of nearshore colonies to converge with their historically slower-growing offshore conspecifics. Bleaching events negatively impacted extension rates in *S. siderea* but not in *P. strigosa*. The more negative trend in linear extension for nearshore versus offshore colonies may arise from ocean warming combined with stronger land-based anthropogenic stressors within nearshore environments. Extrapolating these trends in linear extension into the future suggests that nearshore *P. strigosa* and *S. siderea* will cease growing by years 2110 and 2370, respectively.

## Introduction

Global oceanic change has impacted marine ecosystems worldwide[1], causing range expansions[2], habitat contractions[3], decreased productivity[4], pest outbreaks[5], phase shifts[6], and alterations in ecosystem structure and function[7, 8]. In tropical oceans, sea surface temperatures (SST) have increased by up to 1° C over the past century[9]. As corals inhabiting tropical oceans already live near their thermal maximum[10], even small increases in ocean temperature can have dire consequences for their health and viability. Increased seawater temperature is the primary cause of widespread coral bleaching―a phenomenon in which the obligate coral-algal symbiosis essential for the survival of reef-building corals breaks down, resulting in a white or ‘bleached’ appearance[11]. Mass coral bleaching events have caused significant coral mortality across reef ecosystems globally[12]. These mass bleaching events are of particular concern in the Caribbean Sea, where seawater temperature has increased at higher rates than in other tropical basins[13]. This warming has caused coral cover to decrease by up to 80% in recent decades[14], declines in the structural complexity of local reefs[15], and previously dominant, massive, long-lived coral species to be replaced by smaller short-lived species[16]. If present warming trends continue, bleaching events on Caribbean coral reefs are predicted to increase in both frequency and severity, potentially occurring every two years as soon as 2030[17] and annually by 2040[18]. This increased rate of bleaching, triggered by exposure to more intense, frequent and/or prolonged thermal stress, is predicted to negatively impact rates of coral growth and survival even in more thermally tolerant species.

Coral growth response to temperature has been shown to be parabolic[19]. Moderate increases in temperature below the thermal maximum promote coral growth[19, 20], while temperature increases above the thermal maximum cause coral growth to decline[19]. The duration of corals’ responses to thermal stress can vary based on local factors, as coral growth rates on reefs with higher local stress have been shown to recover to pre-stress growth levels slower than conspecifics from reefs with lower local stress[21]. Nevertheless, more sustained decreases in skeletal extension and calcification have been attributed primarily to ocean warming[20], independent of other local stressors.

Irrespective of cause, declining coral growth rates may increase the incidence of post-settlement mortality in young corals by increasing the duration of exposure to size-specific agents of mortality[19]. As a result, the dominant species of Caribbean coral reefs should continue to shift from fast-growing and structurally complex corals (e.g., *Acropora* sp.) to smaller, fast-growing species[16] (e.g., *Porites* sp.) and larger, slow-growing, domical, stress-tolerant species (e.g., *S. siderea*)[22]. Such shifts in community structure, coupled with decreasing growth rates of surviving corals and increased juvenile mortality, may reduce structural complexity of reefs and decrease rates of gross community calcification. If gross community calcification fails to exceed gross CaCO_3_ dissolution, this will lead to net community dissolution, degradation of the physical reef structure, and collapse of the reef ecosystem that relies upon this structure[19, 22]. Although thermal stress is known to be one of the most negative stressors impacting rates of coral calcification and skeletal extension, coral growth is also impacted by disease, changing ocean chemistry (e.g., ocean acidification), eutrophication, increased sedimentation, food availability, storm activity, and other anthropogenic and non-anthropogenic stressors[19].

Sedimentation and nutrient loading, shown to negatively impact coral skeletal growth parameters[23, 24], are often higher on nearshore reefs than on offshore reefs due to proximity to land—the ultimate source of sediments and nutrients[23, 25]. For example, nearshore massive *Porites* sp. corals on the Great Barrier Reef (GBR) have exhibited decreasing growth rates since 1930, while growth rates in offshore and mid-channel reefs have remained relatively stable[26]. Conversely, *Orbicella annularis* corals exhibited elevated skeletal extension rates in less turbid waters in Jamaica[23], as did *Porites* spp. in Indonesia[24], suggesting that higher water quality supports higher rates of coral growth. However, growth rates of *O. annularis* in Mexico[27] and *Porites* spp. on the GBR[28] are reported to be higher in more turbid waters. Furthermore, in areas of the Florida Keys, nearshore corals exhibited higher growth rates than offshore corals despite their exposure to higher levels of local stress[29].

Elevated and/or increasing growth rates on nearshore reefs with generally lower water quality may be driven by historical exposure to greater temperature variability, which has been shown to confer resilience to corals exposed to anthropogenic thermal stress[30]. Notably, in southern Belize, forereef *S. siderea* corals exhibited declining skeletal extension over the past century, while skeletal extension of nearshore and backreef corals remained relatively stable[31]. Declining skeletal extension in forereef *S. siderea* was correlated with increasing SST, while skeletal extension for backreef and nearshore corals was uncorrelated with SST, suggesting that forereef corals are more vulnerable to thermal stress. The authors attributed this to the fact that forereef corals were historically exposed to less diurnal and seasonal thermal variability, and therefore could be less adapted for anthropogenic warming than backreef and nearshore corals[32].

These differences in historical extension rates in nearshore, backreef, and forereef corals highlight the geographic variability in corals’ response to warming and raise questions about the ultimate driver(s) (e.g., nutrients, sedimentation, and history of thermal exposure) of this variability. Understanding the role that these factors play in corals’ response to ocean warming will provide insight into corals’ ability (or inability) to maintain localized ecosystem function as the oceans continue to warm. This should allow for improved, site-specific management of coral reef ecosystems during this interval of rapid global change.

Here, we investigate the geographic variability of two scleractinian coral species’ response to warming throughout the Mesoamerican Barrier Reef System (MBRS). Century-scale skeletal extension rates were quantified for two abundant and widely distributed massive Caribbean reef-building corals―*Siderastrea siderea* and *Pseudodiploria strigosa*―across numerous nearshore-offshore (i.e., nearshore-backreef-forereef) transects of the Belize MBRS. These transects were selected to represent stress gradients, decreasing from nearshore-to-offshore, because corals in nearshore habitats are exposed to higher summer temperatures, increased thermal variability (diurnal and seasonal), more days per year above the bleaching threshold[33], elevated nutrients (vis-à-vis chlorophyll-a) [33], and greater local anthropogenic stress (e.g., sedimentation, pollution) than offshore corals (backreef, forereef, atolls) due to their proximity to mainland Belize[25, 34].

A total of 134 coral cores were extracted from 19 reef sites across numerous inshore-offshore transects along the entire *ca.* 300 km Belize portion of the MBRS. Colonies of *S. siderea* were sampled from five distinct reef environments (nearshore, backreef, forereef, atoll backreef, atoll forereef) while *P. strigosa* colonies were sampled from two reef environments (nearshore, forereef; Fig. 1). Skeletal extension rates were reconstructed from the thickness of annual high-low density bands identified via X-ray computed tomography (CT). Because skeletal extension rates in both species were strongly linearly correlated with CT-derived calcification rates, the investigation was confined to the skeletal extension data. These data were evaluated for reef-zone differences in annual coral extension rate, slope of coral extension vs. time, and correlation with mass-bleaching events.

**Fig 1:**
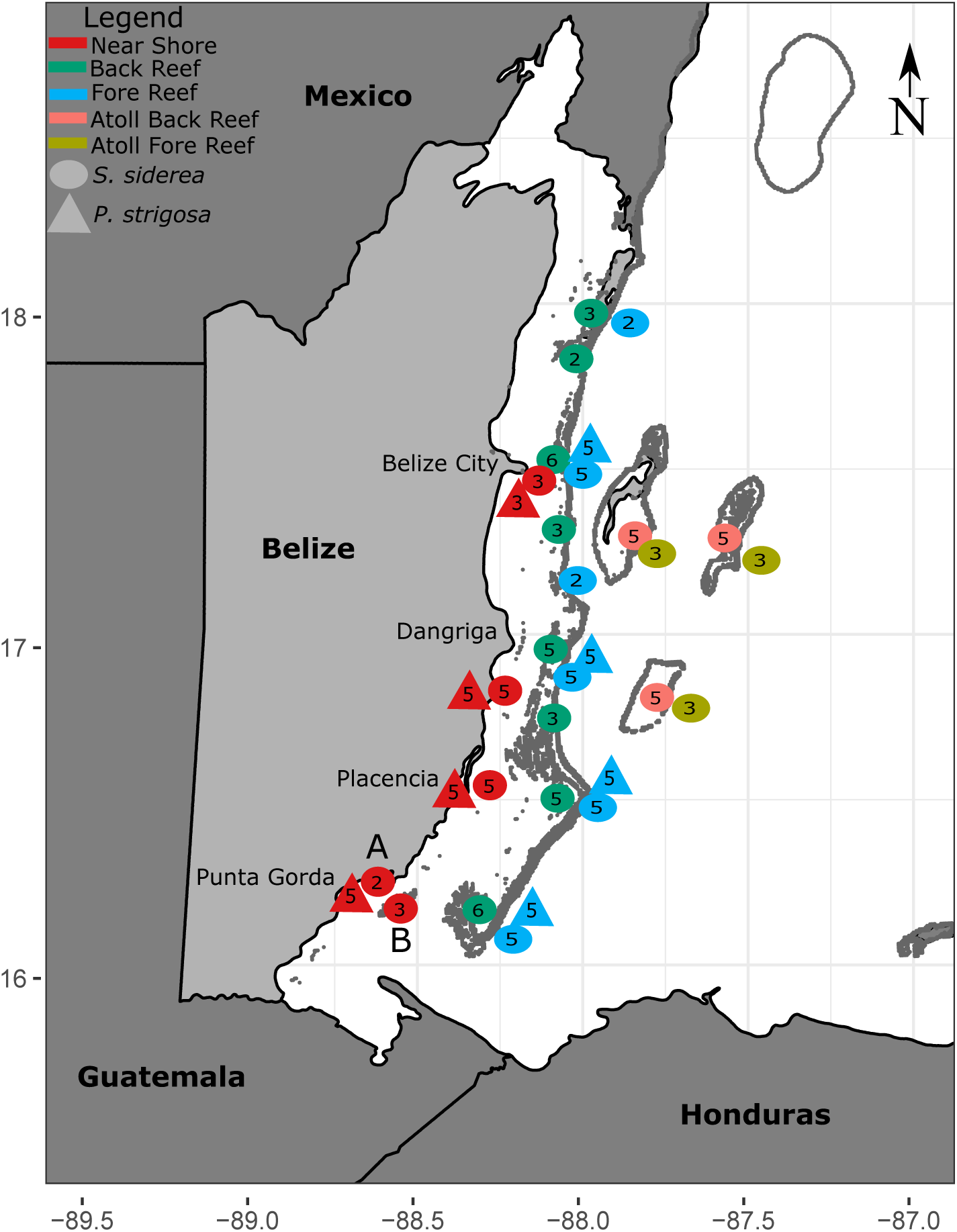
Map of reef sites on the Belize Mesoamerican Barrier Reef where *Sidereastrea siderea* and *Pseudodiploria strigosa* cores were extracted in 2009, 2012, and 2015. Circles and triangles represent core extraction sites for *S. siderea* and *P. strigosa*, respectively. Colors denote reef zone (nearshore = red, backreef = green, forereef = blue, atoll backreef = pink, and atoll forereef = yellow). Numbers denote total cores extracted for a particular species at a specific site.

## Materials and Methods

### Site Description

This research was conducted along the coast of the Belize portion of the Mesoamerican Barrier Reef System (MBRS)—a 1,200 km network of reefs in the western Caribbean sea extending south from the tip of the Yucatan Peninsula in Mexico, traversing the entire coast of Belize and the Atlantic coast of Guatemala, and culminating in the Islas de la Bahia (Bay Islands) off the coast of Honduras (Fig 1).

### Extraction of coral cores

A total of 134 coral cores (93 *S. siderea* and 31 *P. strigosa*) were collected from 19 sites along the Belize MBRS in 2009, 2012, and 2015 (Table S3). All *P. strigosa* cores were collected in 2015. Thirty-seven *S. siderea* cores were collected in 2015, while the remaining 56 *S. siderea* cores were collected in 2009 and 2012. Cores were obtained from five different reef zones (nearshore, backreef, forereef, atoll backreef, atoll forereef) (Fig 1). Backreef, forereef, atoll backreef, and atoll forereef are referred to collectively as offshore reefs. Nearshore coral cores were obtained from within 10 km of the coast of Belize at 4 different latitudes. Backreef and forereef coral cores were obtained on the shoreward and seaward sides of the reef crest, respectively. Corals were transported back to UNC Chapel Hill and CT scanned whole in order to quantify skeletal density, extension, and calcification (see supplementary methods for CT procedures).

### Skeletal density, extension, and calcification

*Siderastrea siderea* and *P. strigosa* are known to deposit one low-density and one high-density growth band per year (seasonally)[35, 36]. Semi-annual density bands were visualized on 8-10 mm thick “slabs” of stacked images (0.6 mm slices) using “mean” projection mode. Mean projection mode utilizes the mean density of each pixel within the 8-10 mm slab, in contrast to min projection mode that uses the minimum density at that pixel within the slice, and maximum projection mode that uses the maximum density[37]. Each annual band pair was demarcated using the “length” tool (ROI drawing tool) in Osirix. Annual linear extension rates for cores were estimated from the thickness of high-density and low-density annual couplets for each core either using the applet RUNNINGCORALGUI (for cores from 2009 and 2012) or manual delineation (see supplementary methoids). Three sets of linear transects were drawn down the length of the cores using the ROI tool in Horos. The linear extension of each seasonal light and dark band was then quantified from the total length of the line tool data in pixels, which was then converted to cm.

Extension, density, and calcification rate were quantified for all corals collected in 2015 (38 *P. strigosa* and 37 *S. siderea*), while only extension was quantified for corals collected in 2009 and 2012 (56 *S. siderea*). Density and calcification rate (calculated from linear extension and density) were not available for cores collected prior to 2015 because the cores were slabbed and sampled for geochemical analysis before they could be CT-scanned with an appropriate density standard. Density of cores collected in 2015 was determined as described above. Corals were oriented with the growth axis parallel to the length of the scanning table to decrease impacts of beam-hardening on density.

### Statistical Analyses

Statistical analyses were performed on individual *S. siderea* and *P. strigosa* core chronologies rather than on a single master chronology for corals from different sub-environments of the reef system [31]. This statistical approach was employed to address the inherent hierarchical nature of coral skeletal growth data. Although all three skeletal growth parameters (skeletal density, extension rate, calcification rate) were quantified for the cores collected in 2015, we focus here on annual skeletal extension because extension is highly correlated with calcification rate (i.e., annual skeletal density does not vary with time; [Fig S1; 19, 20, 38].

Annual skeletal extension rates within a core are inevitably highly correlated across time and therefore are not independent observations, but are approximately independent amongst different cores within the same reef sub-environment. A linear regression of annual skeletal extension with time was achieved by fitting a set of mixed effects models that treated the individual core as a structural variable (Tables 1, S4, supplementary methods). A residual temporal correlation structure was employed to determine if random effects adequately accounted for the correlation over time. To assess the need for random effects, the method of generalized least squares was employed to fit a corresponding set of models with residual correlation structures but without random effects. The use of mixed effects and time series methods to model coral skeletal growth data correctly distinguishes observational units from sampling units, recognizes that sampling variation exists both within and between core time series records, and addresses the temporal autocorrelation structure that is inherently present in such data. The use of mixed effects and time series methods also properly accounts for data imbalance—the fact that some cores provide a longer time series of annual skeletal extension than others. Further information on model testing is available in the supplementary methods.

**Table 1:**
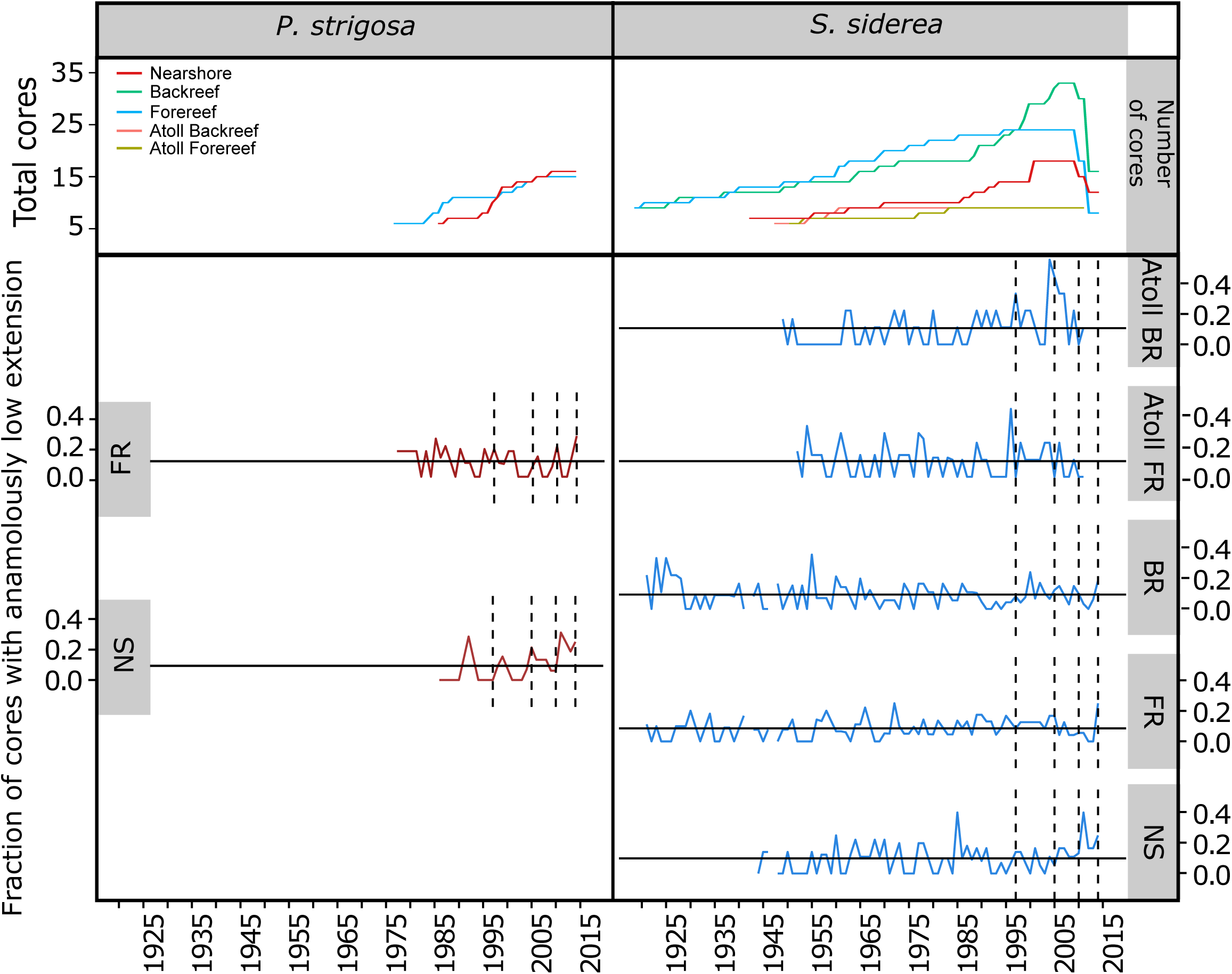
Slope of extension rate by reef zone from linear mixed effects models by species and time scale. Significant *p-values* (*p*<0.05) are in bold and indicate a difference from zero. 95% confidence intervals (CI) that do not overlap indicate significant differences between reef zones (see Fig 2, 3, S2, S3).

## Declining skeletal extension rates for nearshore corals

The slopes of the annual skeletal extension rates vs. time for nearshore *S. siderea* from the late 19^th^ century to present (Fig. 2A, B) and nearshore *P. strigosa* from the mid-20^th^ century to present (Fig. 3A, B) were significantly negative (*p*-values <0.001; Table S1), indicating declining rates of skeletal extension for both coral species on nearshore reefs on the Belize MBRS. In contrast, *S. siderea* and *P. strigosa* colonies from the backreef, forereef, atoll backreef, and atoll forereef (collectively defined as “offshore” because of their >30 km distance from mainland Belize) exhibited relatively stable rates of skeletal extension through time (Fig. 2A, B).

**Fig 2:**
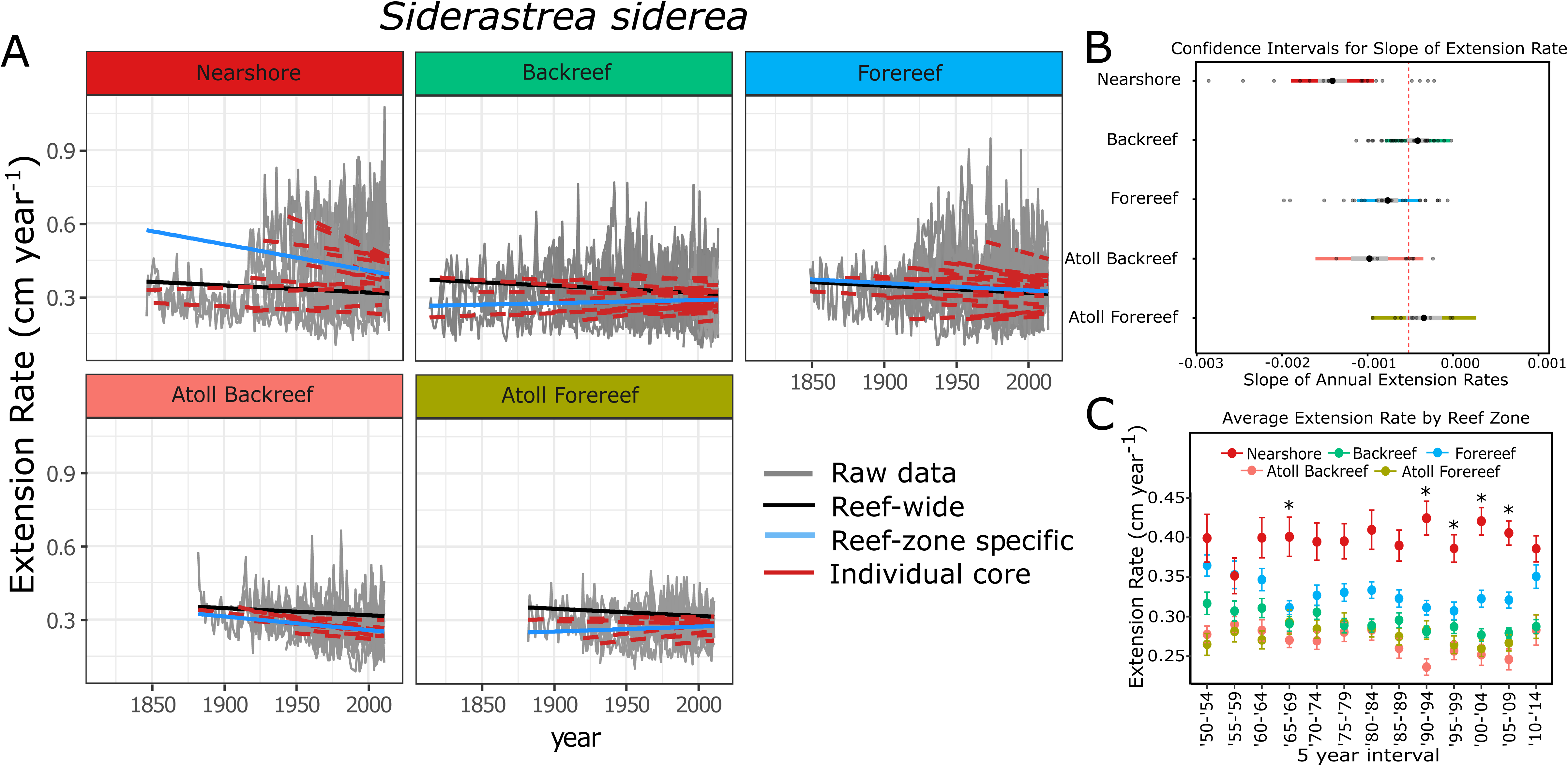
(A) Results of linear model of extension rate (cm year^-1^) for *S. siderea* by reef zone for the 1814-to-present interval. Gray lines are raw extension data, black lines are average linear models of extension for all *S. siderea* cores across all reef zones, blue lines are average linear models of extension for all *S. siderea* cores within each reef zone, and red lines are linear models of extension for individual *S. siderea* cores within reef zones. (B) Slopes of linear models describing extension vs. time for each reef zone, with small points representing individual cores and large points representing average slopes of all cores within a reef zone (gray bars and colored bars are 50% and 95% confidence intervals (CI), respectively, of average slope for each reef zone). Slopes are significantly different from each other if their 95% CI do not overlap. Likewise, slopes are significantly different from zero if their 95% CI do not overlap with the red dashed 0 line. (C) Five-year averages of skeletal extension rate by reef zone ± 1 SE. Asterisks indicate statistically significant differences (p < 0.05) between nearshore and forereef values.

**Fig 3:**
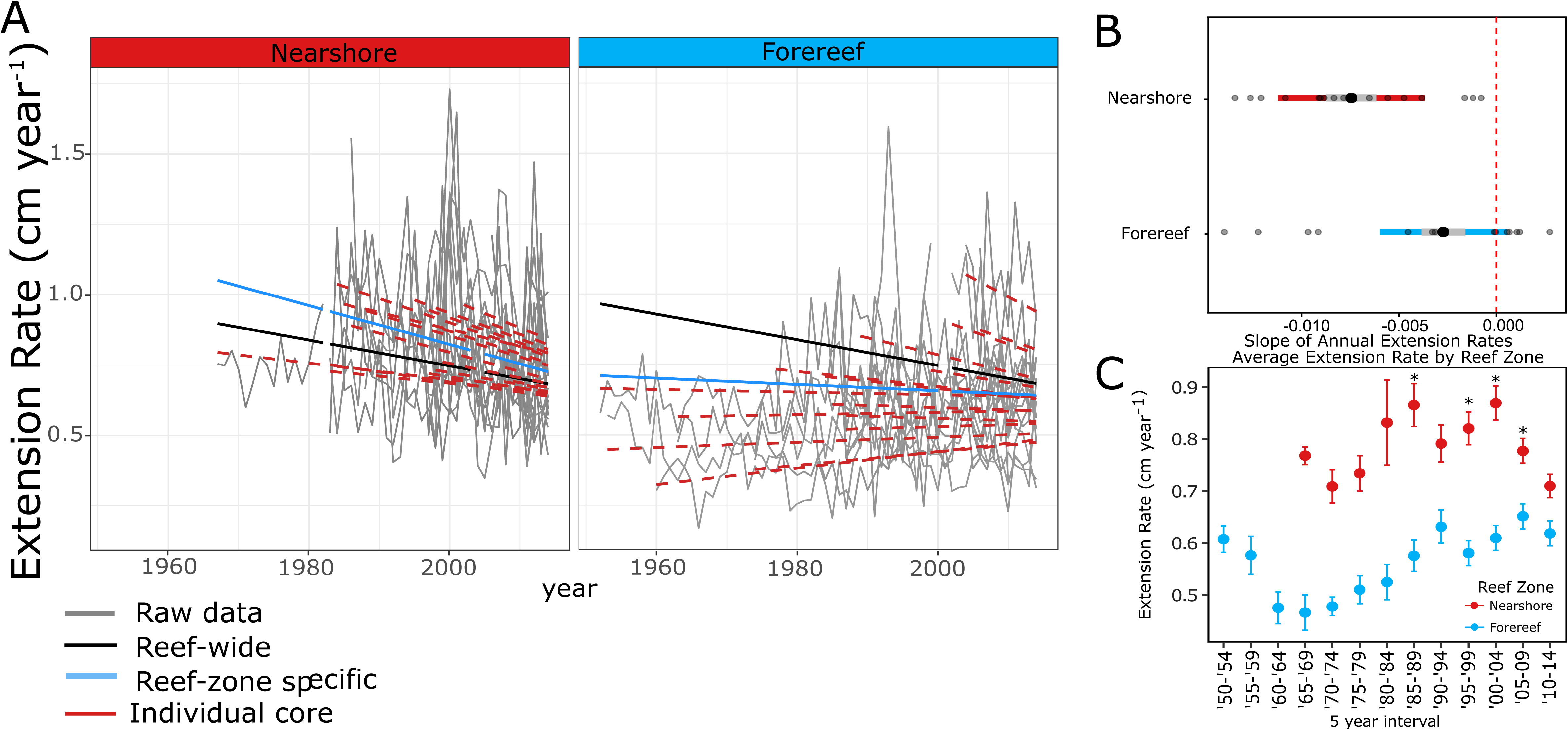
Results of linear model of extension rate (cm year^-1^) for *P. strigosa* by reef zone for the 1950-to-present interval. Gray lines are raw extension data, black lines are average linear models of extension for all *P. strigosa* cores across all reef zones, blue lines are average linear models of extension for all *P. strigosa* cores within each reef zone, and red lines are linear models of extension for individual *S. siderea* cores within reef zones. (B) Slopes of linear models describing extension vs. time for each reef zone, with small points representing individual cores and large points representing average slopes of all cores within a reef zone (gray bars and colored bars are 50% and 95% confidence intervals (CI), respectively, of average slope for each reef zone). Slopes are significantly different from each other if their 95% CI do not overlap. Likewise, slopes are significantly different from zero if their 95% CI do not overlap with the red dashed 0 line. (C) Five-year averages of skeletal extension rate by reef zone ± 1 SE. Asterisks indicate statistically significant differences (p < 0.05) between nearshore and forereef values.

Declining skeletal extension rates for nearshore *S. siderea* and *P. strigosa* corals may be driven by increasing seawater temperatures on nearshore reefs[19, 30], although local stressors such as eutrophication and sedimentation may have also played important roles[39, 40]. Nearshore reefs on the Belize MBRS are exposed to warmer summers and more variable water temperatures than their offshore counterparts, and are subject to greater intervals when temperatures are above the regional bleaching threshold[32, 33]. Additionally, the average SST across all reef zones of the Belize MBRS has increased since 1880 (*p*<0.01; Fig. S2A) and average summer SST across this reef ecosystem has increased by approximately 0.5°C since 1985[32]. However, moderate increases in temperature (below a coral’s thermal optimum) have been shown to increase coral growth rates[19, 41], which may partially explain why nearshore corals exhibit faster growth down-core (i.e., when seawater temperatures were still below the corals’ thermal optimum) than their offshore counterparts in the present study (Fig. 2C; Fig. 3C).

Although temperature increases up to and slightly beyond a coral species’ thermal optimum can increase coral skeletal growth rates[19], temperatures surpassing this thermal optimum by more than a degree have been shown to negatively impact coral growth[19, 42]. This negative impact of elevated temperature on coral skeletal growth rate is driven not only by the magnitude of the warming, but also by its duration[19]. Century-scale declines in skeletal extension rates of nearshore colonies along the Belize MBRS and relatively stability in extension rate of backreef and forereef colonies (Fig. 2; Fig. 3) suggest that a critical threshold of thermal stress (e.g., frequency and/or intensity) may have been exceeded for nearshore *S. siderea* and *P. strigosa* corals, but not for forereef and backreef colonies.

The authors are not aware that a thermal optimum has been established for *P. strigosa*; however, Castillo et al (2014)[41] identified a thermal optimum for *S. siderea* in the range of 28 °C. Furthermore, a regional bleaching threshold (always warmer than a species’ thermal optimum) of 29.7 °C has been identified for various species of corals across the Belize MBRS[43]. Although corals at all sites on the Belize MBRS are exposed to temperatures above this threshold each year, nearshore reefs on the Belize MBRS are exposed to between 54 and 78 days per year above the bleaching threshold of 29.7 °C, with sustained intervals above the bleaching threshold lasting up to 7.5 consecutive days. In contrast, offshore reef sites experience only 20 to 40 days above the bleaching threshold annually, with sustained intervals above the bleaching threshold lasting fewer than 4.8 consecutive days[33].

These observations, combined with the observation that extension rates on nearshore reefs have been declining over the past century while extension rates for offshore reefs have been relatively stable over the past century, suggest that the thermal threshold for temperature-related declines in coral growth lies somewhere between temperatures at these two reef locations. Alternatively, other environmental factors, such as ocean acidification, eutrophication, and/or sedimentation, may be driving the negative growth trends observed for nearshore reefs, but not for offshore reefs, of the Belize MBRS[19].

Previous work has demonstrated that poor water quality impairs coral growth rates on nearshore reefs[26]. Specifically, coral calcification rates on nearshore reefs of the GBR are declining on multi-decadal timescales, while calcification rates on offshore reefs are increasing. This declining growth on nearshore reefs is attributed to the impacts of wet season river discharge of sediment and nutrients, a trend that is exacerbated by warming [26]. In the present study, it is possible that increasing nutrient and sediment loading[25, 44], coupled with increasing water temperatures and duration of time when water temperatures exceed the species’ bleaching threshold, are responsible for the decline in skeletal extension rates observed on nearshore reefs of the Belize MBRS.

In Belize, human population densities have increased 39% in coastal cities and agricultural land area has quadrupled since the mid-21^st^ century (Fig. S2B, C). It is therefore likely that runoff and eutrophication in nearshore environments of the MBRS have also increased over time[25, 45]. This increase in runoff and eutrophication should impact water quality more negatively at nearshore reefs than at offshore reefs that are further from the pollution source[25]. If temperature and eutrophication continue to increase, nearshore coral growth rates should continue to decline—with offshore corals potentially following suit as these stressors impact more distal portions of the Belize MBRS. Although there is metagenomic evidence that nearshore *S. siderea* and possibly *P. strigosa* have begun acclimatizing to these elevated temperatures[46], the observation that skeletal extension rates have continued declining for both species up to present time in nearshore reefs of the MBRS indicate that such acclimatization within nearshore corals is insufficient for maintaining stable rates of skeletal growth amidst the deteriorating environmental conditions of nearshore environments.

Further evidence of the combined effects of warming and local stress on nearshore coral skeletal extension is observed at the southernmost nearshore site in the present study. At this location, *S. siderea* cores were extracted from two nearshore reefs, one nearer to the coast at Sheepshead Caye (6 km from mainland Belize; point A in Fig. 1) and one further from the coast at Snake Cayes (13 km from mainland Belize; point B in Fig. 1). The cores collected from Snake Cayes (farther from mainland; NS14, NS15, NS16) exhibited the lowest skeletal extension rates of any nearshore core analyzed in the present study (Table S7; Table S8; Fig. S4), but did not exhibit declining skeletal extension rates with time[31], while those from Sheepshead Caye (closer to mainland) did exhibit declining skeletal extension rates with time.

Corals from Sheepshead Caye were exposed to warmer and more variable temperatures, on average, than corals from Snake Cayes[33], and since corals from Sheepshead Caye are nearer to the mainland, it is likely that they are subject to greater eutrophication and sedimentation than corals from Snake Cayes (Fig. S2). The impacts of higher temperature, greater thermal variability, increased sedimentation, and/or eutrophication appear to have combined to cause declining skeletal extension rates at Sheepshead Caye, while the relatively lower temperature, thermal variability, sedimentation, and/or eutrophication, owing to Snake Cayes’ greater distance from the mainland, were insufficient to cause coral growth at that location to significantly decline with time. These shore-distance gradients in skeletal extension trends within nearshore reefs of the southern MBRS recapitulate the regional scale, nearshore-offshore MBRS trend in coral skeletal extension rates, in which extension rates of nearshore corals have declined with time, while extension rates of offshore (backreef, forereef, atoll) corals have exhibited relative stability with time.

## Extension rates of nearshore corals have decreased to the level of offshore conspecifics

Nearshore *S. siderea* and *P. strigosa* exhibited higher skeletal extension rates than offshore conspecifics from at least 1990 to 2009 (Table S2; Table 3; Fig. 2C, 3C; *p*-values <0.001). This trend is visually apparent as far back as 1965, but decreasing sample size further back in time diminishes the statistical significance of this relationship (Table S2; Table 3; Fig. 2C, 3C). After 2009, however, skeletal extension rates of nearshore *S. siderea* and *P. strigosa* converge with those of their offshore conspecifics (*p-*values: 0.986 and 0.186, respectively; Table S2; Table 3; Fig. 2C; Fig. 3C) owing to the decline in skeletal extension rates for the nearshore corals.

The results from the present study suggest that warmer and more nutrient-rich nearshore reef environments historically supported higher skeletal extension rates than offshore reef environments (Table S2; Table 3; Fig. 2C; Fig. 3C; Fig. S4). On the Belize MBRS, nearshore reefs have historically experienced higher temperatures than offshore reefs[33] and since warming below a species’ thermal optimum can increase coral growth rates[19], it is not surprising that nearshore corals historically (pre-2010) exhibited higher skeletal extension rates than offshore corals inhabiting historically cooler waters. Furthermore, proximity to shore (i.e., source of nutrients and sediments) dictates that nutrient levels and sediment load are likely higher on nearshore reefs than on offshore reefs (Fig. S2)[25, 47].

Although decreased light availability from increased turbidity (i.e., elevated suspended sediment, algal blooms) can inhibit coral growth[39] and nutrient enrichment [and subsequent altering of nitrogen (N):phosphorus (P) ratio][40, 48] can increase bleaching susceptibility and lead to decreased growth rates[23, 26], some coral species, including *S. siderea* and *P. strigosa*, metabolize N from ingested sediments and particulates [49, 50]. This N may augment coral nutrition during intervals of increased sedimentation and eutrophication, potentially mitigating some of the negative impacts of these processes. Additionally, exposure to increased N and P, when coupled with heterotrophic feeding, has been shown to enhance coral calcification, even when the corals are exposed to thermal stress[51]. A combination of enhanced nutrition and elevated temperatures (below the bleaching threshold) may have been responsible for nearshore corals growing faster than offshore corals before 2010 (Fig. 2C; Fig. 3C).

However, the recent convergence of extension rates for nearshore and offshore colonies of *S. siderea* and *P. strigosa* (Fig. 2) suggests that the advantage that nearshore corals appear to have historically had over offshore corals has now been lost, perhaps due to the intense warming, eutrophication, and sedimentation targeting nearshore environments over recent decades (Figs. 2, 3, S2; Table S2; Table 3). These declining trends in skeletal extension may also have significant impacts on the geomorphology of nearshore reef environments, as slowing growth can result in reef-scale flattening and a loss of structural complexity that may impact the ecological function of nearshore reefs[15] and may ultimately impair their ability to keep pace with rising sea level.

## Recent bleaching events differentially impact corals across reef environments

Mass coral bleaching was documented in the Caribbean in 1997-1998, 2005, 2010, and 2014-2016 (see methods and Donner et al., 2017[52]). The skeletal extension data from the present study was evaluated to determine whether recent mass bleaching events in the Caribbean Sea impacted coral skeletal extension within each reef zone of the Belize MBRS[53]. Overall, skeletal extension was significantly lower during bleaching years than during non-bleaching years for *S. siderea* (*p*<0.001; Table S6), but not for *P. strigosa*, although there were some bleaching years in which *P. strigosa* exhibited significantly lower extension than during non-bleaching years (Table S5; Fig. S3). These relationships between bleaching and extension did not vary significantly by reef zone for either species (Table S6). However, skeletal extension was anomalously low for *S. siderea* on the forereef and backreef of the atolls during the 1997-1998 bleaching event and on the backreef of the atolls following the 2005 bleaching event (Table S5; Fig. S3), for nearshore *S. siderea* and *P. strigosa* following the 2010 bleaching event (Table S5; Fig. S3), and for nearshore *S. siderea* and forereef corals of both species during the 2014 bleaching event (Table S5; Fig. S3). Notably, anomalously low skeletal extension rates were also observed for some non-bleaching years in both species (e.g., in 1985 for nearshore *S. siderea* and in 1992 for nearshore *P. strigosa*; Table S5; Fig. S3), potentially due to other stressors (e.g. storms, human activity, or sedimentation[12, 19]) or unreported/small-scale bleaching.

Anomalously low skeletal extension rates for both nearshore and forereef conspecifics in the same year were observed only in *S. siderea* in 2014, indicating that the impact of this bleaching event on growth of this species was more widespread than past bleaching events, possibly resulting from the cumulative impacts of increased temperatures, bleaching, and/or local stressors in the preceding years. The 2010 bleaching event correlated with low extension for both species in the nearshore reef zone, but not in the other reef zones, demonstrating that the impact of individual bleaching events on coral skeletal extension varied across reef zones, even though the general relationship between all bleaching events and coral extension did not vary significantly across reef zones.

Although single mass bleaching events were correlated with low rates of skeletal extension within some reef zones, no single bleaching event was correlated with low rates of skeletal extension across all reef zones, underscoring the variability in how individual bleaching events impact skeletal extension across coral species and reef environments. Therefore, the declining skeletal extension rates observed on nearshore reefs of the Belize MBRS cannot be confidently attributed to the increasing frequency of mass bleaching events in recent years. Instead, the steady nature of the decline in skeletal extension of the investigated species in nearshore reef environments suggests that it has been caused by the comparably steady increase in seawater temperatures over the same interval, which is also the root cause of the bleaching events themselves. Nevertheless, the increasing frequency of the bleaching events may indeed be exacerbating the deleterious impacts of steady anthropogenic warming on skeletal extension rates in these nearshore reef environments.

## Results predict that nearshore colonies of *P. strigosa* will cease growing by year 2110

Extrapolating from historical growth trends, skeletal extension of nearshore *S. siderea* corals of the MBRS is expected to decline by 23% by year 2100 and to cease entirely by year 2374 ± 17, while skeletal extension of nearshore *P. strigosa* of the MBRS is expected to decline by 89% by year 2100 and to cease entirely by year 2110 ± 34. Although both species are considered stress-tolerant[54], substantial differences in their historical trends in skeletal extension suggest that *S. siderea* is more stress-tolerant than *P. strigosa*. Less stress-tolerant corals would naturally be expected to suffer even more depressed extension and to cease growing earlier. Coral reefs are predicted to transition to a state of net dissolution by the end of the present century due to the impacts of ocean acidification on carbonate sediment dissolution, assuming little to no decline in coral calcification[55]. Our results suggest that coral calcification on nearshore reefs along the Belize MBRS will decline drastically over the next century, even in the most stress-tolerant species, suggesting that nearshore reef platforms (i.e., living corals and algae plus non-living reef frameworks and sediments) of the MBRS will experience net dissolution well before the end of the century. The resulting degradation of the three-dimensional reef structure and collapse of the associated reef ecosystem will lead to species extirpation and/or extinction, decreasing coral diversity and evenness, reef-flattening, and loss of reef complexity and habitat on nearshore reefs of the MBRS[15, 22].

These predicted declines in coral growth assume that the temporal trends in coral extension observed over the cored interval can be linearly extrapolated into the future, which is predicated on the assumptions that the primary coral stressors (e.g., warming, acidification, eutrophication, sedimentation, pollution) will continue changing at the same rate and that corals’ responses to these stressors will be linear. However, continued improvement of local water quality and reduction in global CO_2_ emissions (if achieved) have the potential to mitigate some of these projected growth decreases. For example, emissions scenarios lower than or on par with the commitments of the Paris Agreement have been projected to potentially increase or at least maintain stable growth rates for Bermudan corals[56]. Conversely, further deterioration of water quality and/or acceleration of warming and acidification beyond rates observed over the cored interval and/or development of synergistic impacts amongst stressors would accelerate future declines in coral extension.

## Declining skeletal extension in nearshore corals foretells deterioration of entire MBRS

The results of the present study reveal a clear difference in historical growth trends between nearshore and offshore corals of the Belize MBRS. Nearshore *S. siderea* and *P. strigosa* historically exhibited higher skeletal extension rates compared to their offshore conspecifics (Fig. 2; Fig. 3). This higher growth of nearshore corals was likely driven by historically warmer temperatures—favorable to the extent that they were below the corals’ thermal optimum—and lower local environmental stress [25, 44] (Fig. S2), although other factors may have played a role. However, extension rates of nearshore *S. siderea* and *P. strigosa* have now declined to levels similar to their historically slower growing offshore conspecifics owing to seawater temperatures more frequently exceeding the corals’ thermal optima and from higher local environmental stress in nearshore environments.

Although skeletal extension trends of offshore corals have exhibited relative stability over the observed interval, the decline in extension rate of nearshore colonies that are presently experiencing sustained thermal stress beyond their thermal optimum may foretell future declines in the growth of offshore colonies once their thermal optima are more consistently exceeded.

Declines in extension of nearshore colonies of both species do not reliably correlate with mass bleaching events—suggesting that the long-term decline in nearshore coral extension cannot be unequivocally attributed to the increasing frequency of mass bleaching events. Instead, long-term increases in seawater temperature and local stressors (e.g., eutrophication and sedimentation), which are typically more pronounced in nearshore environments owing to their mainland proximity, are the more likely drivers of the observed decline in nearshore coral growth. Any advantage historically conferred to corals by inhabiting the nearshore environment, vis-à-vis thermal acclimation and/or increased heterotrophic uptake of N and/or C in particle-rich nearshore waters, has now been lost.

Furthermore, continued declines in coral growth could lead to complete stoppage of growth by year 2110 for nearshore *P. strigosa* and by year 2370 for nearshore *S. siderea*. Such a scenario would cause baseline dissolution rates of coral skeletons, which have also been shown to increase with warming[57], to exceed rates of gross coral calcification. This, coupled with increasing carbonate sediment dissolution[55], would result in the net reef dissolution (i.e., gross dissolution > gross calcification) and eventual collapse and disappearance of nearshore reef structures[58].

Although nearshore corals historically exhibit higher rates of growth than offshore corals, rapid deterioration of environmental conditions in nearshore environments has caused growth rates of nearshore corals to approach those of their offshore conspecifics. Such declines in these and other reef species are reducing the biodiversity, structure, and ecosystem function of the Belize MBRS[15, 22]. Continued protection and management of these reefs should include monitoring land use to limit increases in sedimentation and eutrophication of reefs (particularly nearshore reefs), as well as local, regional, and global action to reduce CO_2_ emissions and stabilize global temperatures and ocean pH. The rapid and persistent decline in skeletal extension of two species of nearshore corals underscores the urgency of this action, which might afford corals of the Belize MBRS sufficient time to acclimatize to and, hopefully, survive this interval of rapid climate and oceanic change.

## Data availability

Data will be available at Dryad when this paper is published doi:10.5061/dryad.jn680jd

## Authors’ contributions

JHB designed the study, carried out the research, carried out the statistical analysis, and wrote the manuscript; JBR conceived of the study, provided resources, and helped draft the manuscript; JPR helped carry out the research, helped design the statistical analysis, and carried out the research; TAC designed the study and contributed to statistical analysis; HEA coordinated the field component of the study; IW helped carry out the study; KDC conceived of and coordinated the study. All authors gave final approval for publication.

## Acknowledgements

We thank K. Horvath, K. Cobleigh, C. Hines, A. Rash, Garbutt’s Marine, and the Toledo Institute for Development and the Environment (TIDE) for field assistance, A. Foguel for development of the *RunningCoral* program for quantifying coral extension, Wake Radiology and UNC Biomedical Research Imaging Center (BRIC) for assistance with coral CT scanning, and C. Campbell, C. Anderson, D. DeLeener, H. Knight, J. Boulton, J. Townsend, M. Roycroft, M. Mendoza, P. Armstrong, and V. Eastman for lab assistance. We also thank the Belize Fisheries Department for issuance of the relevant research permits and for their continued support of our research efforts on the Belize MBRS. This research was funded by a Rufford Foundation grant to J. Baumann, NOAA grants NA11OAR431016 and NA13OAR4310186 to J. Ries and K. Castillo, and National Science Foundation grants to K. Castillo (OCE 1459522) and J. Ries (OCE 1437371).

## References

1. Walther G.-R., Post E., Convey P., Menzel A., Parmesan C., Beebee T.J.C., Fromentin J.-M., Hoegh-Guldberg O., Bairlein F. 2002 Ecological responses to recent climate change. Nature 416(389–395).

2. Elmhagen B., Kindberg J., Hellström P., Angerbjörn A. 2015 A boreal invasion in response to climate change? Range shifts and community effects in the borderland between forest and tundra. AMBIO 44(1), 39–50. (doi:10.1007/s13280-014-0606-8).

3. Smale D.A., Wernberg T. 2013 Extreme climatic event drives range contraction of a habitat-forming species. Proceedings of the Royal Society B: Biological Sciences 280(1754). (doi:10.1098/rspb.2012.2829).

4. O’Reilly C.M., Alin S.R., Plisnier P.-D., Cohen A.S., McKee B.A. 2003 Climate change decreases aquatic ecosystem productivity of Lake Tanganyika, Africa. Nature 424(6950), 766–768. (doi:http://www.nature.com/nature/journal/v424/n6950/suppinfo/nature01833_S1.html).

5. Kurz W.A., Dymond C.C., Stinson G., Rampley G.J., Neilson E.T., Carroll A.L., Ebata T., Safranyik L. 2008 Mountain pine beetle and forest carbon feedback to climate change. Nature 452(7190), 987–990. (doi:http://www.nature.com/nature/journal/v452/n7190/suppinfo/nature06777_S1.html).

6. Connell S.D., Russell B.D. 2010 The direct effects of increasing CO_2_ and temperature on non-calcifying organisms: increasing the potential for phase shifts in kelp forests. Proceedings of the Royal Society B: Biological Sciences. (doi:10.1098/rspb.2009.2069).

7. Hoegh-Guldberg O., Bruno J.F. 2010 The Impact of Climate Change on the World’s Marine Ecosystems. Science 328(5985), 1523–1528. (doi:10.1126/science.1189930).

8. Knowlton N. 2001 The future of coral reefs. Proceedings of the National Academy of Sciences 98(10), 5419–5425.

9. Deser C., Phillips A.S., Alexander M.A. 2010 Twentieth century tropical sea surface temperature trends revisited. Geophysical Research Letters 37(10).

10. Glynn P.W. 1993 Coral reef bleaching: ecological perspectives. Coral Reefs 12, 1–17.

11. Jokiel P.L., Coles S.L. 1990 Response of Hawaiian and other Indo-Pacific reef corals to elevated temperature. Coral Reefs 8, 155–162.

12. Hughes T.P., Kerry J.T., Álvarez-Noriega M., Álvarez-Romero J.G., Anderson K.D., Baird A.H., Babcock R.C., Beger M., Bellwood D.R., Berkelmans R. 2017 Global warming and recurrent mass bleaching of corals. Nature 543(7645), 373–377.

13. Chollett I., Mumby P.J., Müller-Karger F.E., Hu C. 2012 Physical environments of the Caribbean Sea. Limnology and Oceanography 57(4), 1233–1244.

14. Gardner T.A., Cote I.M., Gill J.A., Grant A., Watkinson A.R. 2003 Long-term region-wide declines in Caribbean corals. Science 301, 958–960.

15. Alvarez-Filip L., Dulvy N.K., Gill J.A., Côté I.M., Watkinson A.R. 2009 Flattening of Caribbean coral reefs: region-wide declines in architectural complexity. Proceedings of the Royal Society of London B: Biological Sciences, rspb20090339.

16. Green D., Edmunds P., Carpenter R. 2008 Increasing relative abundance of Porites astreoides on Caribbean reefs mediated by an overall decline in coral cover. Marine Ecology Progress Series 359, 1–10. (doi:10.3354/meps07454).

17. Donner S.D., Knutson T.R., Oppenheimer M. 2007 Model-based assessment of the role of human-induced climate change in the 2005 Caribbean coral bleaching event. Proceedings of the National Academy of Science 104(13), 5483–5488.

18. Van Hooidonk R., Maynard J.A., Liu Y., Lee S.K. 2015 Downscaled projections of Caribbean coral bleaching that can inform conservation planning. Global Change Biology.

19. Pratchett M.S., Anderson K.D., Hoogenboom M.O., Widman E., Baird A.H., Pandolfi J.M., Edmunds P.J., Lough J.M. 2015 Spatial, temporal and taxonomic variation in coral growth—implications for the structure and function of coral reef ecosystems. Oceanography and Marine Biology: An Annual Review 53, 215–296.

20. Lough J., Barnes D. 2000 Environmental controls on growth of the massive coral Porites. Journal of Experimental Marine Biology and Ecology 245(2), 225–243.

21. Carilli J.E., Norris R.D., Black B.A., Walsh S.M., McField M. 2009 Local stressors reduce coral resilience to bleaching. Plos ONE 4(7), e6324.

22. Alvarez-Filip L., Carricart-Ganivet J.P., Horta-Puga G., Iglesias-Prieto R. 2013 Shifts in coral-assemblage composition do not ensure persistence of reef functionality. Scientific reports 3.

23. Dodge R.E., Aller R.C., Thomson J. 1974 Coral growth related to resuspension of bottom sediments. Nature 247, 574–576.

24. Tomascik T. 1990 Growth rates of two morphotypes of Montastrea annularis along a eutrophication gradient, Barbados, WI. Marine Pollution Bulletin 21(8), 376–381.

25. Heyman W.D., Kjerfve B. 1999 Hydrological and oceanographic considerations for integrated coastal zone management in southern Belize. Environmental Management 24(2), 229–245.

26. D’Olivo J., McCulloch M., Judd K. 2013 Long-term records of coral calcification across the central Great Barrier Reef: assessing the impacts of river runoff and climate change. Coral Reefs 32(4), 999–1012.

27. Carricart-Ganivet J.P., Merino M. 2001 Growth responses of the reef-building coral Montastraea annularis along a gradient of continental influence in the southern Gulf of Mexico. Bulletin of Marine Science 68(1), 133–146.

28. Lough J., Barnes D., Devereux M., Tobin B., Tobin S. 1999 Variability in growth characteristics of massive Porites on the Great Barrier Reef. CRC Reef Research Centre Technical Report (28), 95.

29. Lirman D., Fong P. 2007 Is proximity to land-based sources of coral stressors an appropriate measure of risk to coral reefs? An example from the Florida Reef Tract. Marine Pollution Bulletin 54(6), 779–791.

30. Carilli J., Donner S.D., Hartmann A.C. 2012 Historical temperature variability affects coral response to heat stress. Plos ONE 7(3), e34418.

31. Castillo K., Ries J., Weiss J. 2011 Declining coral skeletal extension for forereef colonies of Siderastrea siderea on the Mesoamerican Barrier Reef System, southern Belize. PLoS ONE 6(2), e14615.

32. Castillo K.D., Ries J.B., Weiss J.M., Lima F.P. 2012 Decline of forereef corals in response to recent warming linked to history of thermal exposure. Nature Climate Change 2(10), 756–760.

33. Baumann J.H., Townsend J.E., Courtney T.A., Aichelman H.E., Davies S.W., Lima F.P., Castillo K.D. 2016 Temperature Regimes Impact Coral Assemblages along Environmental Gradients on Lagoonal Reefs in Belize. Plos ONE 11(9), e0162098. (doi:10.1371/journal.pone.0162098).

34. Carilli J.E., Norris R.D., Black B., Walsh S.M., McFIELD M. 2010 Century-scale records of coral growth rates indicate that local stressors reduce coral thermal tolerance threshold. Global Change Biology 16(4), 1247–1257.

35. Guzman H.M., Tudhope A.W. 1998 Seasonal variation in skeletal extension rate and stable isotopic (^13^C/^12^C and ^18^O/^16^O) composition in response to several environmental variables in the Caribbean reef coral Siderastrea siderea. Marine Ecology Progress Series 166, 109–118.

36. Helmle K.P., Dodge R.E., Ketcham R. 2000 Skeletal architecture and density banding in Diploria strigosa by X-ray computed tomography.

37. Carilli J.E., Hartmann A.C., Heron S.F., Pandolfi J.M., Cobb K., Sayani H., Dunbar R., Sandin S.A. 2017 Porites coral response to an oceanographic and human impact gradient in the Line Islands. Limnology and Oceanography.

38. Lough J., Barnes D. 1997 Several centuries of variation in skeletal extension, density and calcification in massive Porites colonies from the Great Barrier Reef: A proxy for seawater temperature and a background of variability against which to identify unnatural change. Journal of Experimental Marine Biology and Ecology 211(1), 29–67.

39. Fabricius K.E. 2005 Effects of terrestrial runoff on the ecology of corals and coral reefs: review and synthesis. Marine Pollution Bulletin 50(2), 125–146.

40. Wiedenmann J., D’Angelo C., Smith E.G., Hunt A.N., Legiret F.-E., Postle A.D., Achterberg E.P. 2013 Nutrient enrichment can increase the susceptibility of reef corals to bleaching. Nature Climate Change 3(2), 160–164.

41. Castillo K.D., Ries J.B., Bruno J.F., Westfield I.T. 2014 The reef-building coral Siderastrea siderea exhibits parabolic responses to ocean acidification and warming. Proceedings of the Royal Society of London B: Biological Sciences 281(1797), 20141856.

42. Lough J.M., Cantin N.E. 2014 Perspectives on massive coral growth rates in a changing ocean. The Biological Bulletin 226(3), 187–202.

43. Aronson R.B., Precht W.F., Toscano M.A., Koltes K.H. 2002 The 1998 bleaching event and its aftermath on a coral reef in Belize. Marine Biology xxx.

44. Thattai D., Kjerfve B., Heyman W. 2003 Hydrometeorology and variability of water discharge and sediment load in the inner Gulf of Honduras, western Caribbean. Journal of Hydrometeorology 4(6), 985–995.

45. Carilli J.E., Prouty N.G., Hughen K.A., Norris R.D. 2009 Century-scale records of land-based activities recorded in Mesoamerican coral cores. Marine Pollution Bulletin 58(12), 1835–1842.

46. Davies S.W., Ries J.B., Marchetti A., Granzotti R., Castillo K.D. 2017 Symbiodinium functional diversity and clade specificity under global change stressors. bioRxiv. (doi:10.1101/190413).

47. Chérubin L., Kuchinke C., Paris C. 2008 Ocean circulation and terrestrial runoff dynamics in the Mesoamerican region from spectral optimization of SeaWiFS data and a high resolution simulation. Coral Reefs 27(3), 503–519.

48. Rosset S., Wiedenmann J., Reed A.J., D’Angelo C. 2017 Phosphate deficiency promotes coral bleaching and is reflected by the ultrastructure of symbiotic dinoflagellates. Marine Pollution Bulletin 118(1), 180–187.

49. Mills M.M., Sebens K.P. 2004 Ingestion and assimilation of nitrogen from benthic sediments by three species of coral. Marine Biology 145, 1097–1106.

50. Mills M.M., Lipschultz F., Sebens K.P. 2004 Particulate matter ingestion and associated nitrogen uptake by four species of scleractinian corals. Coral Reefs 23(3), 311–323.

51. Ezzat L., Towle E., Irisson J.O., Langdon C., Ferrier-Pagès C. 2015 The relationship between heterotrophic feeding and inorganic nutrient availability in the scleractinian coral T. reniformis under a short-term temperature increase. Limnology and Oceanography.

52. Donner S.D., Rickbeil G.J., Heron S.F. 2017 A new, high-resolution global mass coral bleaching database. Plos ONE 12(4), e0175490.

53. Andréfouët S., Berkelmans R., Odriozola L., Done T., Oliver J., Müller-Karger F. 2002 Choosing the appropriate spatial resolution for monitoring coral bleaching events using remote sensing. Coral Reefs 21(2), 147–154. (doi:10.1007/s00338-002-0233-x).

54. Darling E.S., Alvarez-Filip L., Oliver T.A., McClanahan T.R., Côté I.M. 2012 Evaluating life-history strategies of reef corals from species traits. Ecology Letters 15(12), 1378–1386.

55. Eyre B.D., Cyronak T., Drupp P., De Carlo E.H., Sachs J.P., Andersson A.J. 2018 Coral reefs will transition to net dissolving before end of century. Science 359(6378), 908–911.

56. Courtney T.A., Lebrato M., Bates N.R., Collins A., de Putron S.J., Garley R., Johnson R., Molinero J.-C., Noyes T.J., Sabine C.L. 2017 Environmental controls on modern scleractinian coral and reef-scale calcification. Science advances 3(11), e1701356.

57. Ries J.B., Ghazaleh M.N., Connolly B., Westfield I., Castillo K.D. 2016 Impacts of seawater saturation state (ΩA=0.4–4.6) and temperature (10, 25°C) on the dissolution kinetics of whole-shell biogenic carbonates. Geochimica et Cosmochimica Acta 192, 318–337. (doi:https://doi.org/10.1016/j.gca.2016.07.001).

58. Eyre B.D., Andersson A.J., Cyronak T. 2014 Benthic coral reef calcium carbonate dissolution in an acidifying ocean. Nature Climate Change 4(11), 969.

